# Evaluating the Safety of Intravenous Injection of Nitric Oxide, Magnesium, and Hydrogen Nano Bubble Solution in Sprague Dawley Rats

**DOI:** 10.1101/2025.06.16.659826

**Authors:** Dody Novrial, Nor Sri Inayati, Nur Signa Aini Gumilas, Dhadhang Wahyu Kurniawan, Sutiman Bambang Sumitro

## Abstract

Nanobubble technology exhibits outstanding stability and permeability and has been extensively developed in pharmaceuticals, dentistry, and medicine. The Hydrogen Nano Bubble (HNB) solution containing NO and magnesium is projected to have an even greater therapeutic effect than has ever been investigated. An experimental study using Sprague-Dawley rats was conducted to assess the intravenous injection’s safety. Each rat received a single injection into the tail vein at a dose of 2 mL. On the other hand, aquades were injected into the rats in the control group. Observations were made every 30 minutes for the first four hours, then every four hours for the first twenty-four hours, and once daily for the next fourteen days. The LD_50_, urea-creatinine level, SGOT/SGPT level, and organ histology, including liver, kidney, heart, lung, and spleen, were evaluated. The solution appears safe for intravenous administration, as there were no adverse side effects during the trial, and all values were still within the normal range. Additional investigation is required to assess the long-term harmful impact.

## Introduction

The therapeutic benefits of high-pressure hydrogen (H2) in treating albino mice with squamous cell carcinoma were first reported in 1975 (Dole et al., 1975). Since then, studies on the gas have progressed and demonstrated considerable promise as an anti-tumor, anti-inflammatory, and antioxidant agent (Iuchi et al., 2016; Sim et al., 2020; Watanabe et al., 2017). Due to the poor accessibility of hydrogen exposure into the body, H2’s application as a therapeutic option remains limited, despite its widespread acceptance as an antioxidant reagent (M. Yang et al., 2020). The poor solubility of H2 in water reduces the effectiveness of oral therapy using hydrogen-rich water (Iida et al., 2016)

Nanobubble technology has been widely developed for applications in medicine, dentistry, and pharmaceuticals, demonstrating excellent stability and permeability (Kato et al., 2020; Xiao & Miwa, 2021). Hydrogen nano bubble (HNB) technology is a cutting-edge innovation that harnesses the power of hydrogen in nanoscale bubbles. These tiny bubbles have unique properties that make them highly effective in various applications, from energy storage to medical treatments (Lohse, 2018). This technology makes it possible to distribute hydrogen to various systems with greater stability and efficiency by encasing the gas in tiny bubbles (Li et al., 2022). The bubbles are perfect for targeted medication administration in medicine because of their small size, making it easy for them to pass through cell membranes (Hansen et al., 2023).

Nitric oxide (NO) is a simple gas made up of two atoms: oxygen (O) and nitrogen (N). Colorless and odorless, NO dissolves readily in both organic and water-based solvents. The NO synthase (NOS) enzyme helps cells in various species, including bacteria, fungi, plants, and animals, create NO as a bioproduct (Nowaczyk et al., 2021). Numerous studies have been published on using NO donors for diagnosis and treatment. Microbubbles laden with NO can improve vasodilation, decrease platelet aggregation, and lessen thrombus formation and inflammatory cell infiltration. This ability is highly beneficial in treating and preventing myocardial infarction and stroke (Huang et al., 2023; Liao et al., 2020; F. Yang et al., 2016).

The mineral Magnesium (Mg^2+^) is present in food, dietary supplements, pharmaceuticals, and the body itself. More than 300 enzyme systems that control various bodily biochemical processes, including protein synthesis, muscle and nerve function, blood glucose control, and blood pressure regulation, depend on Mg^2+^ as a cofactor (Bertinato et al., 2015). From minor symptoms like exhaustion, nausea, and vomiting to more serious ones like hypocalcemia and hypokalemia, Mg^2+^ shortage can induce a range of symptoms. Mg^2+^ deficiency can raise the incidence of migraines, osteoporosis, diabetes, hypertension, and cardiovascular disorders over the long run (Al Alawi et al., 2018).

Overall, HNB technology shows great promise in revolutionizing multiple industries with its versatile and powerful capabilities. It is expected that the solution of HNB with NO and magnesium will have an even greater therapeutic effect. So in this study, we evaluated the acute toxicity potential of that combination.

## Methods

This study was conducted in the research laboratory of the Faculty of Medicine, Jenderal Soedirman University, from August 2024 to January 2025. This research has received ethical approval from the health research ethics committee of the Faculty of Medicine, Universitas Jenderal Soedirman, with ref number: 065/KEPK/PE/VIII/2024.

Sprague Dawley strain white female rats weighing between 150 and 250 grams and aged 8 to 12 weeks were the study’s subjects. Rats must be in optimal wellness, have normal anatomy, have never given birth, and not be pregnant to be included. Their weight must also be distributed evenly among all groups, with variations not to exceed 20% of the test animals’ average body weight. All animals were kept in maintenance cages in a well-ventilated space with temperatures between 25 and 30 degrees Celsius. The lighting was regulated to the day-night cycle. Daily intake with ad libitum aquadest and a typical laboratory-produced rat diet.

Following seven days of quarantine, they were divided into three treatment groups and one control group, each with six rats.

The Indonesia Molecule Institute provided the test material, which came in the form of nanobubbles. The test materials were administered to the treatment groups with the following compositions: NO 5ml + Mg 3ml + Oxygen 3ml + HNB 15ml (A); NO 7.5ml + Mg 3ml + Oxygen 3ml + HNB 12.5ml (B); and NO 10ml + Mg 3ml + Oxygen 3ml + HNB 10ml (C), each in 500ml of NaCl infusion fluid. Each rat received a single intravenous injection into the tail vein at a dose of 2 ml. On the other hand, aquades were injected into the rats in the control group (K). Following test preparation, observations were made every 30 minutes for the first four hours, then every four hours for the first twenty-four hours, and once daily for the next fourteen days.

### Median Lethal Dose (LD_50_)

A median lethal dose (LD50) is observed in this study. LD_50_ is a single dosage of a drug that, according to statistical estimates, will kill 50% of test subjects. LD_50_ is calculated using the Thompson and Weil method in acute toxicity studies (Weil, 1952).

### Blood Sampling

Blood sampling was performed to assess SGOT/SGPT, urea, and creatinine levels through the retro-orbital vein before treatment (I) and after 14 days of post-treatment observation (II).

### Histological Examination

After the observation period was completed, the rats were terminated immediately. The liver, kidney, heart, lung, and spleen were taken for subsequent routine histopathological examination using hematoxylin-eosin staining.

In this study, observations were made on liver histology with a scoring method as follows: Score 1: The liver appears normal; Score 2: There is hydropic/ fatty degeneration/ necrosis focused in one area; Score 3: There is hydropic/ fatty/ necrotic degeneration focused in several areas; Score 4: There is hydropic/ fatty degeneration/ necrosis throughout the area/ field of view. The assessment was conducted on 10 fields of view per sample at a magnification of 200x.

The observation of kidney histology used the scoring method as follows:

Score 1: The kidneys appear normal; Score 2: Lesions such as nuclear necrosis, tubular degeneration, and proximal tubules dilation were found in several fields of view; Score 3: Lesions such as nuclear necrosis, tubular degeneration, and proximal tubules dilation were found in each field of view The assessment was conducted on 10 fields of view per sample at a magnification of 200x.

Observation of heart histology was conducted using the following scoring method:

Score 1: The heart appears normal; Score 2: Mild lesions (inflammation/ fibrosis/ vacuolization <3 foci); Score 3: Moderate lesion (inflammation/ fibrosis/ vacuolization >3 foci); Score 4: Severe lesion (diffuse inflammation/ fibrosis/ vacuolization). The assessment is conducted on 10 fields of view per sample at a magnification of 200x.

The following grading system was used to observe lung histology: Score 1: Normal lung appearance; Score 2: Alveolar wall thickening without lung structural damage; Score 3: Alveolar wall thickening or fibrosis combined with lung architectural damage; Score 4: Significant fibrosis that resembles honeycomb; Score 5: Complete lung injury. Ten fields of view for each sample were evaluated at a magnification of 200x.

In this study, observations of the spleen histology were also conducted using a scoring method as follows: Score 1: Spleen appears normal; Score 2: Focal bleeding/necrosis (mild); Score 3: Multifocal bleeding/necrosis (moderate); Score 4: Diffuse hemorrhage/necrosis (severe). The assessment is conducted on 10 fields of view per sample at 200x magnification.

### Statistical Analysis

Continuous data are presented as mean ± SD, while categorical data are presented as numbers (percentages). The Kruskal-Wallis test, followed by the Mann-Whitney post hoc test, was used in statistical analysis. A p-value < 0.05 indicates a significant difference in the variable’s mean.

## Results and Discussion

### Observation of the animals

According to study findings, there were no appreciable changes in the rats’ behavior, activity, or physical state. During the study period, no test animals died, and none of them suffered from seizures, tremors, or diarrhea. This suggests that the test animals’ physical activity did not decline due to the combination of NO, Mg, and HNB being administered to them.

Measurements of body weight were taken at regular intervals beginning with the acclimatization phase, throughout treatment, one day post-treatment, and then every five days until day fourteen post-treatment. According to the observation results, there has been a consistent daily increase in body weight.

The average body weight of the rats was in the range of 181 grams at the beginning of the study; however, after NO, Mg, and HNB combination was administered, the rats’ body weight increased steadily until it reached the range of 204–207 grams at the end of the experiment (Figure 1). Gaining weight could be a sign that the test substance is not having the harmful effects that lead to weight loss, which is frequently the case when exposed to dangerous compounds. Conversely, weight gain can suggest that the test substance has growth-promoting or appetite-boosting effects on test animals (Chapman et al., 2013; van Berlo et al., 2022).

**Figure 1:**
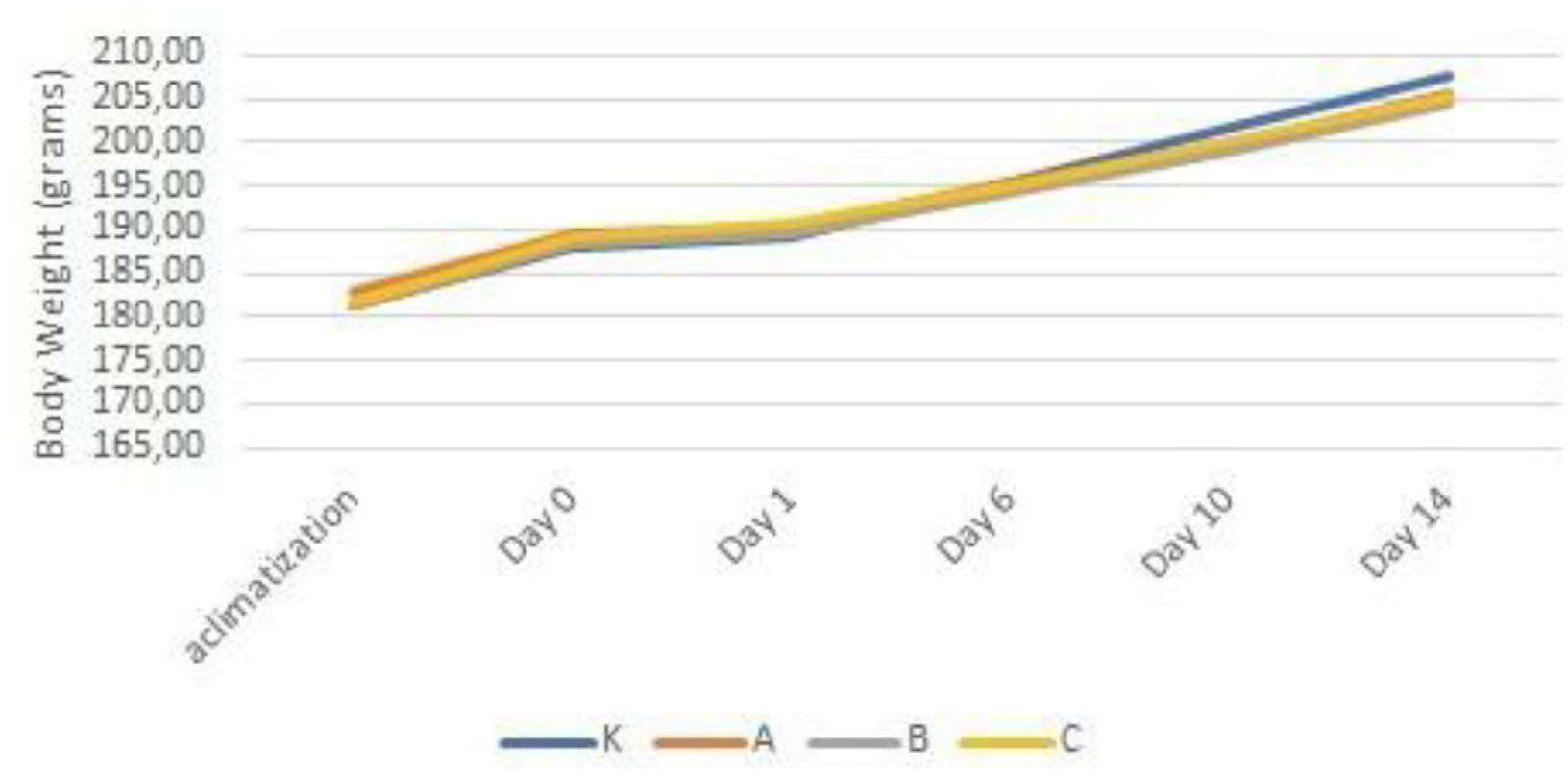
Body weight graph of the rats during the experimental period

During the 14-day observation period, no deaths were found in the test animals across all dosage groups. This suggests that the range of doses tested does not exhibit sufficient toxicity to achieve the LD_50_. Since this LD_50_ cannot be determined, it can be considered that the test substance has low toxicity or that the maximum dose administered to the test animals is insufficient to kill half of the test animal population (Erhirhie et al., 2018). As a result, it may be concluded that the dosage is still largely safe to use as a test dose.

### SGOT-SGPT, Urea, and Creatinine Analysis

The levels of the enzymes serum glutamic pyruvic transaminase (SGPT) and serum glutamic oxaloacetic transaminase (SGOT) were measured to assess liver function. There were significant differences between the SGOT II, SGPT I, and SGPT II levels among the groups, however, there were no significant differences between the SGOT I levels, as shown in Table 1.

**Table 1.** Bivariate analysis of SGOT-SGPT, urea, and creatinine levels between groups.

| Groups | SGOT (U/L) |  |  |  | SGPT (U/L) |  |  |  |
| --- | --- | --- | --- | --- | --- | --- | --- | --- |
|  | I |  | II |  | I |  | II |  |
|  | (Mean±SD) | *p* | (Mean±SD) | *p* | (Mean±SD) | *p* | (Mean±SD) | *p* |
| K | 36.98±0.19 | 0.07 | 37.54±0.32 | 0.02* | 18.13±0.16 | 0.02* | 18.29±0.16 | 0.01* |
| A | 37.38±0.22 |  | 37.79±0.26 |  | 18.29±0.20 |  | 18.61±0.27 |  |
| B | 37.46±0.29 |  | 38.03±0.32 |  | 18.61±0.10 |  | 19.18±0.17 |  |
| C | 37.95±0.23 |  | 38.92±0.15 |  | 18.93±0.22 |  | 19.50±0.23 |  |
| Groups | Urea (mg/dL) |  |  |  | Creatinine (mg/dL) |  |  |  |
|  | I |  | II |  | I |  | II |  |
|  | (Mean±SD) | *p* | (Mean±SD) | *p* | (Mean±SD) | *p* | (Mean±SD) | *p* |
| K | 10.05±0.67 | 0.004* | 10.41±0.15 | 0.01* | 0.77±0.01 | 0.012* | 0.78±0.007 | 0.32 |
| A | 10.69±0.67 |  | 10.88±0.09 |  | 0.79±0.004 |  | 0.81±0.01 |  |
| B | 10.85±0.15 |  | 11.22±0.20 |  | 0.79±0.02 |  | 0.81±0.01 |  |
| C | 11.06±0.23 |  | 11.56±0.21 |  | 0.85±0.002 |  | 0.81±0.02 |  |

\*statistically significant results

Urea and creatinine levels were also analyzed to assess kidney function in each group. Urea and creatinine are important indicators in assessing the kidneys’ ability to filter metabolic waste from the blood. The increase in the levels of these two parameters generally indicates a disturbance in kidney function or nephron damage. Table 1 shows that there were significant differences in urea I and II levels and creatinine I levels between groups, whereas there were no significant differences in creatinine levels II between groups.

The post hoc test findings for SGOT II levels showed that treatment group C had considerably greater levels than treatment groups A, B, and the control group (K), as shown in Table 2. In contrast, treatment groups B and C had considerably greater SGPT I levels than the control group (K). The SGPT II levels in treatment groups B and C were noticeably greater than in the control group. While not substantially different from the SGPT II levels in treatment group B, the SGPT II levels in treatment group C were significantly higher than those in treatment group A.

**Table 2.** Post hoc analysis of SGOT-SGPT, urea, and creatinine levels.

| Groups | *p-value* |  |  |  |  |  |
| --- | --- | --- | --- | --- | --- | --- |
|  | SGOT II | SGPT I | SGPT II | Urea I | Urea II | Creatinine I |
| K-A | 0.68 | 0.49 | 0.25 | 0.003* | 0.039* | 0.057 |
| K-B | 0.36 | 0.03* | 0.01* | 0.005* | 0.015* | 0.684 |
| K-C | 0.01* | 0.01* | 0.01* | 0.005* | 0.008* | 0.003* |
| A-B | 0.68 | 0.21 | 0.10 | 0.309 | 0.141 | 1.000 |
| A-C | 0.01* | 0.05 | 0.03* | 0.242 | 0.033* | 0.003* |
| B-C | 0.04* | 0.21 | 0.26 | 0.514 | 0.222 | 0.096 |

\*statistically significant results

These findings demonstrate how the test substance affects increases in SGOT and SGPT, which are indicators for assessing liver function, as well as the rise in NO and fall in GNH levels during test preparation. All rat groups’ SGOT and SGPT levels, however, remained within the normal range by the end of the trial (normal SGOT levels for white rats are 45.7-80.8 U/L; normal SGPT levels are 17.5-30.2 U/L) (Cathy Johnson Delaney, 2008)

As shown in Table 2, treatment groups A, B, and C had significantly higher urea levels than the control group (K), according to the results of the post hoc tests for urea I and II levels. Additionally, urea II levels in treatment group C were substantially greater than in treatment group A. In contrast, treatment group C had considerably higher values for creatinine I levels than treatment group A and the control group (K), which was possibly caused by individual variation.

These results illustrate the influence of the test substance on the increase in urea and creatinine as markers for kidney function evaluation, along with the increase in NO levels and the decrease in HNB levels in the test preparation. Nevertheless, overall, by the end of the study, the levels of urea and creatinine in rats across all groups remained within the normal range (normal values for white rat urea 10-26.4 mg/dL; creatinine 0.5-0.83 mg/dL)(Arifianto et al., 2020; Zhang & Parikh, 2019).

### Histological Examination

Bivariate analysis of organ histology is presented in Table 3. Significant differences between groups were found in the liver and kidney histologic scores. In the liver, scores of 3 and 4 out of 4 were more commonly found in the treatment groups B and C (Figure 2). On the other hand, the frequency distribution of kidney histology is evenly divided between scores 1 and 2, with all rats in treatment group C having a score of 2 out of 3 (Figure 3). Scoring of heart, lung, and spleen histology did not show significant differences between groups. This analysis is consistent with the observed histological findings, with no significant damage in all treatment groups (Figures 4-6).

**Table 3.**
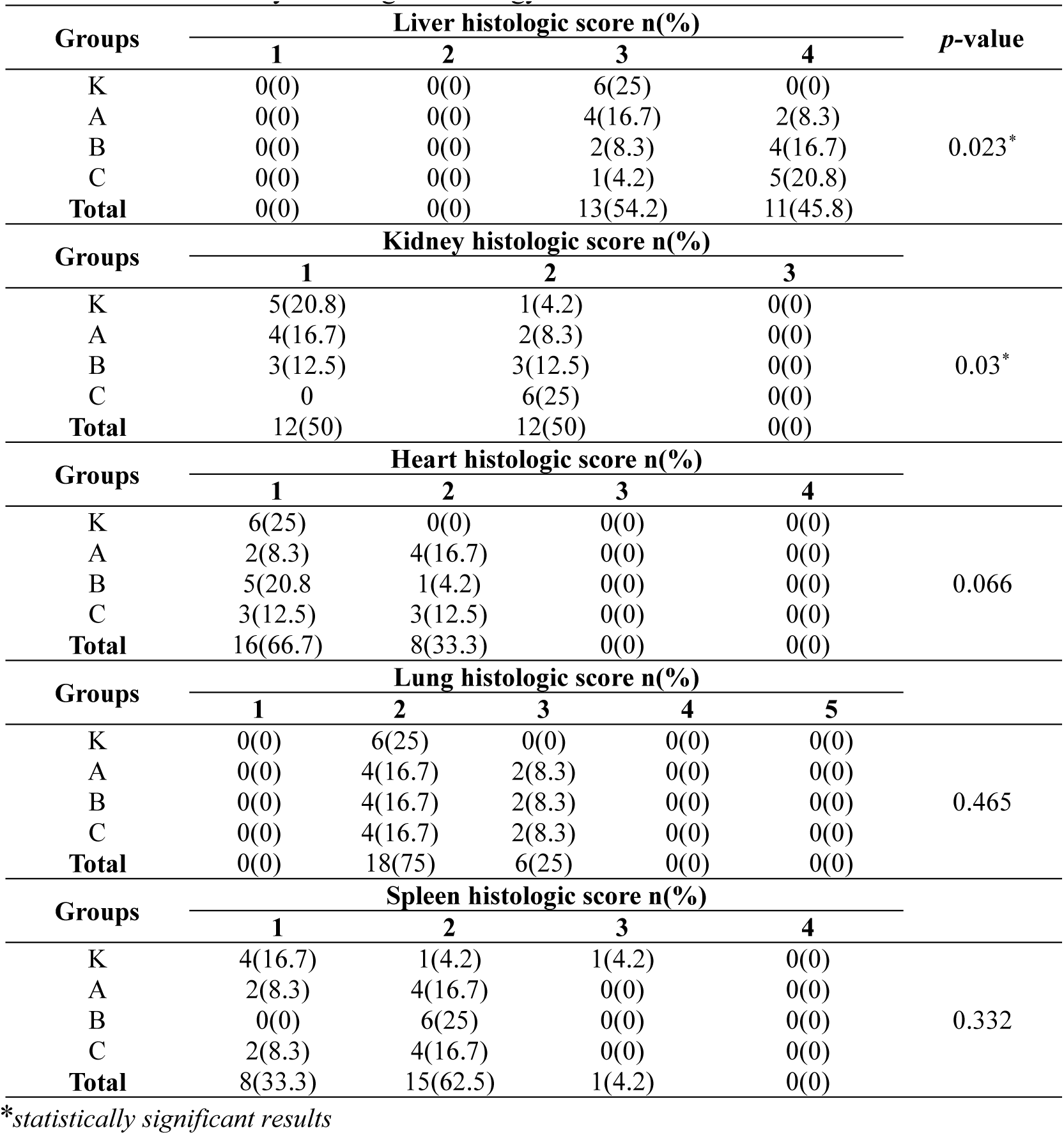
Bivariate analysis of organ histology.

**Figure 2.**
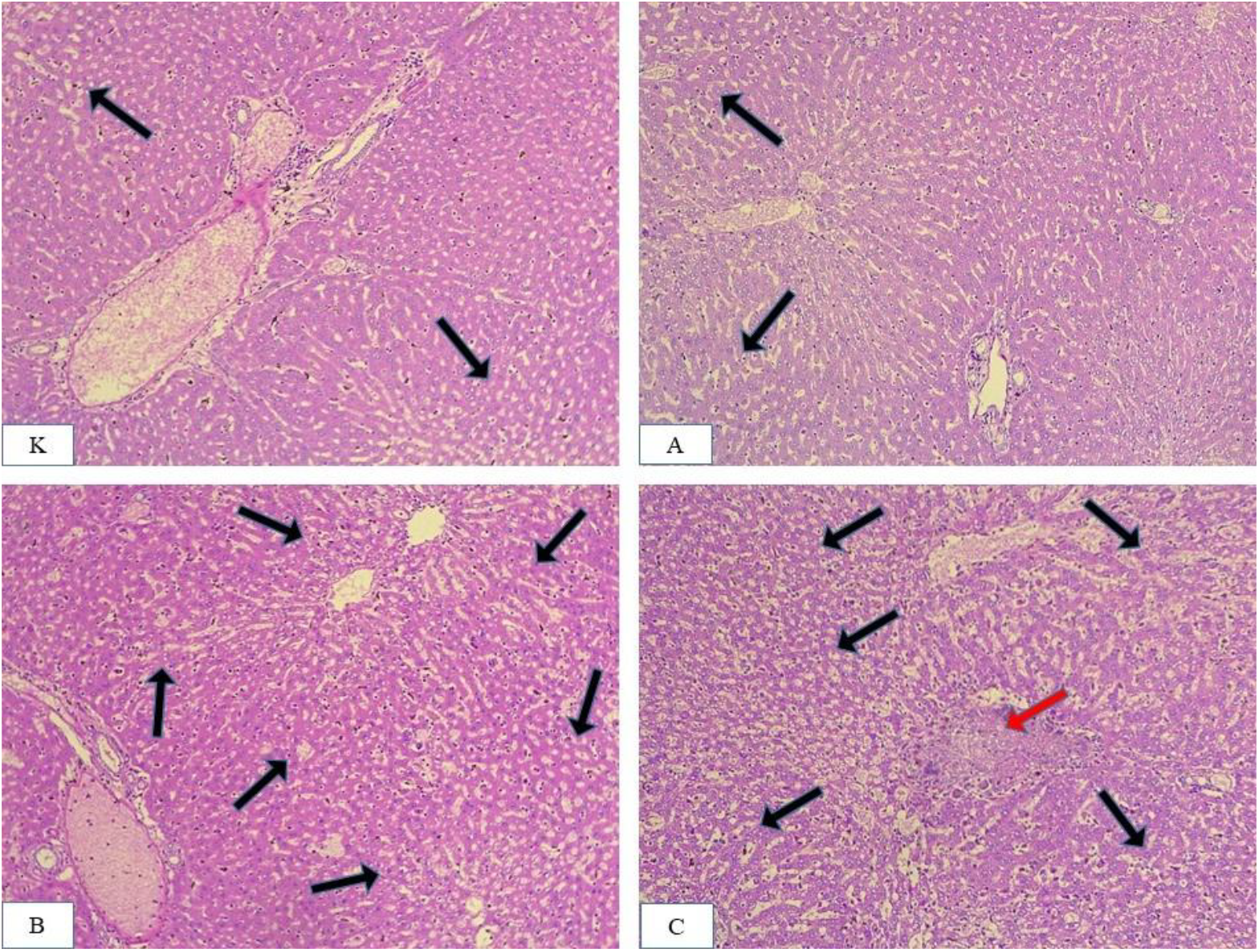
Liver histology. Multifocal fatty degeneration is observed in groups K and A (black arrow). Diffuse fatty degeneration is observed in groups B and C (black arrow), accompanied by necrosis (red arrow). HE 200x

**Figure 3.**
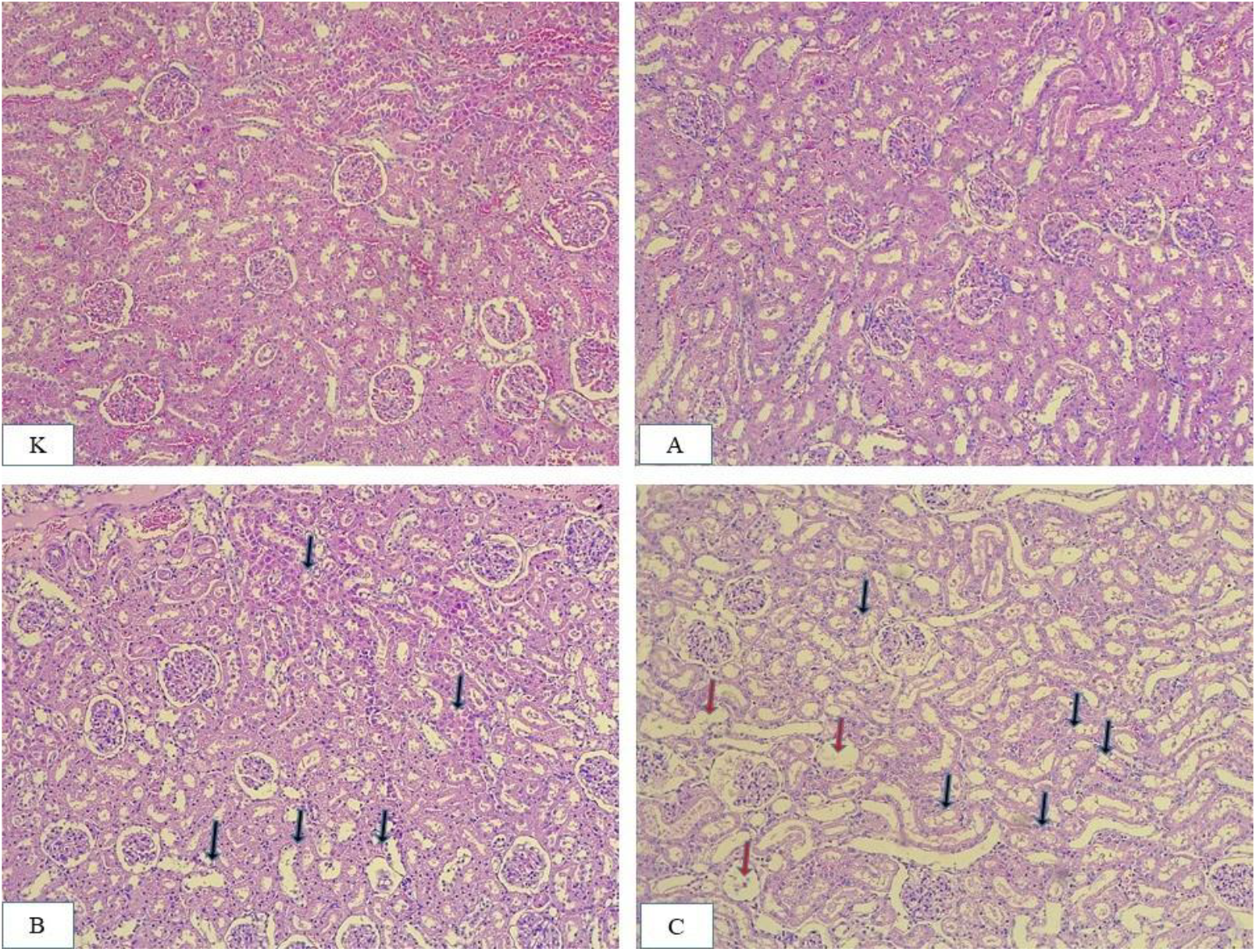
Histology of normal kidneys in groups K and A. Multifocal tubular degeneration (black arrow) is observed in groups B and C. Tubule dilation is observed in group C (red arrow). HE 200x.

**Figure 4.**
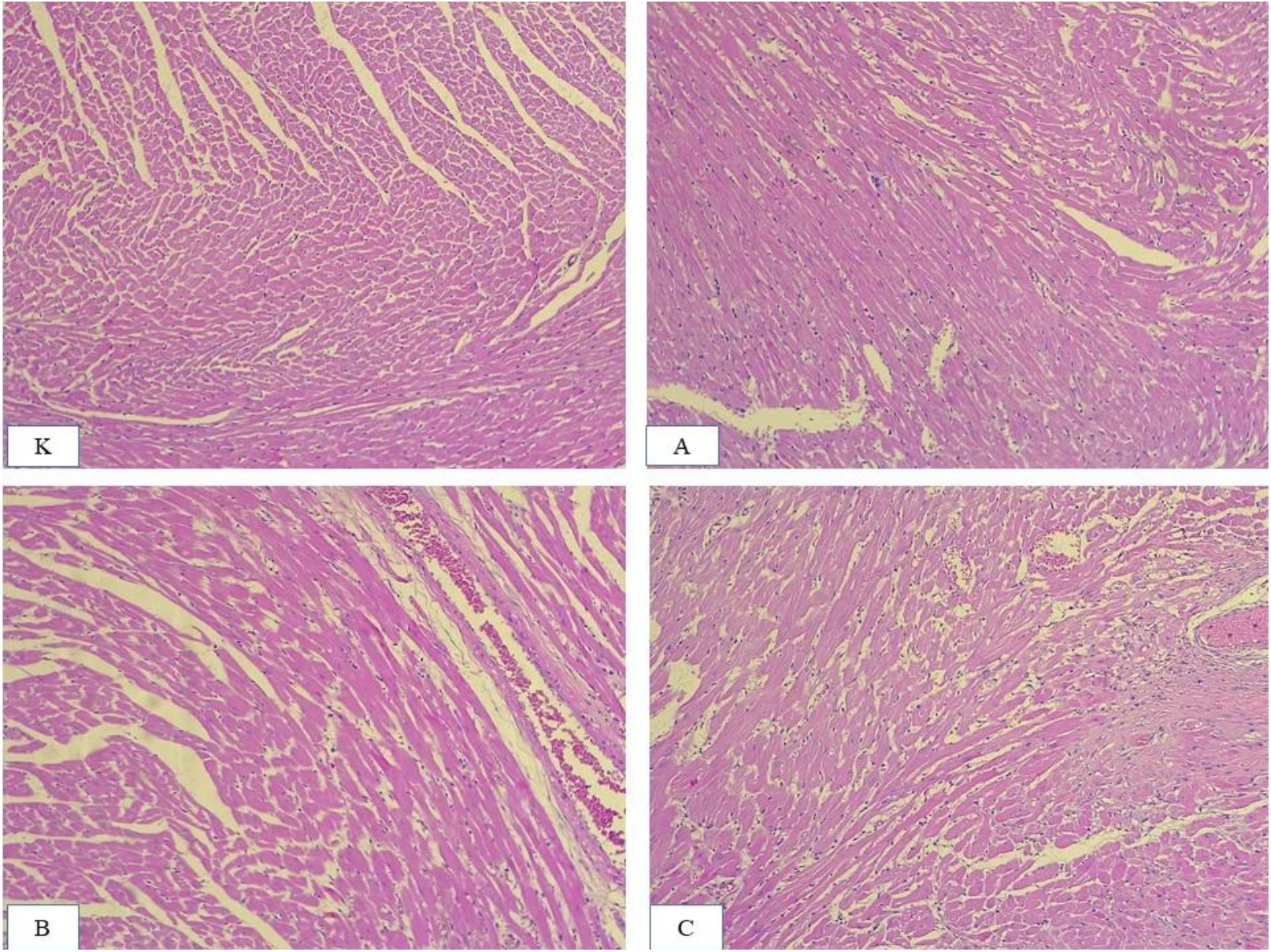
Heart histology appears normal in the K group, with slight inflammatory lesions/light fibrosis observed in the treatment group. HE 200x.

**Figure 5.**
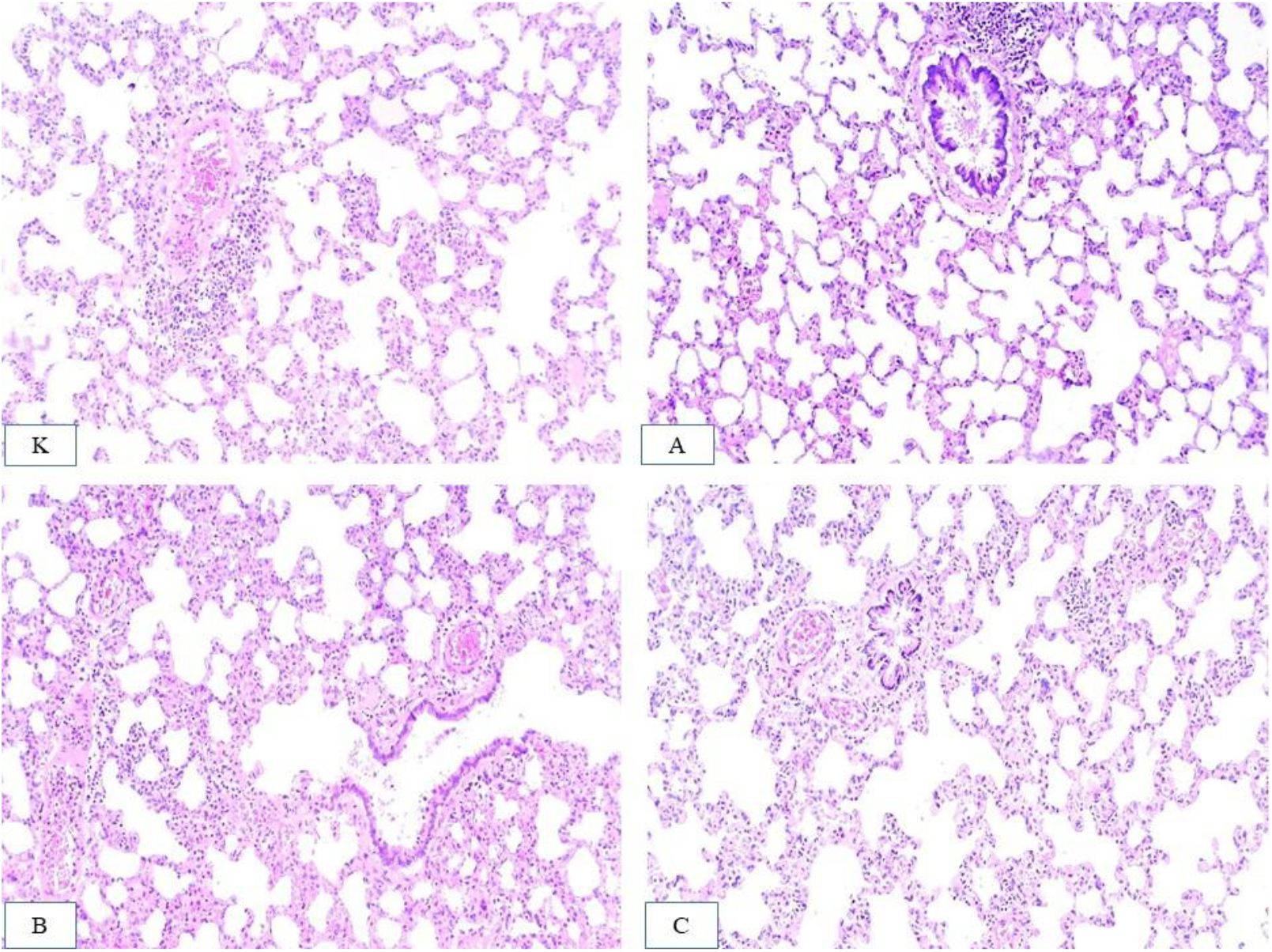
Lung histology in all groups shows mild inflammation without damaging the lung structure. HE 200x.

**Figure 6.**
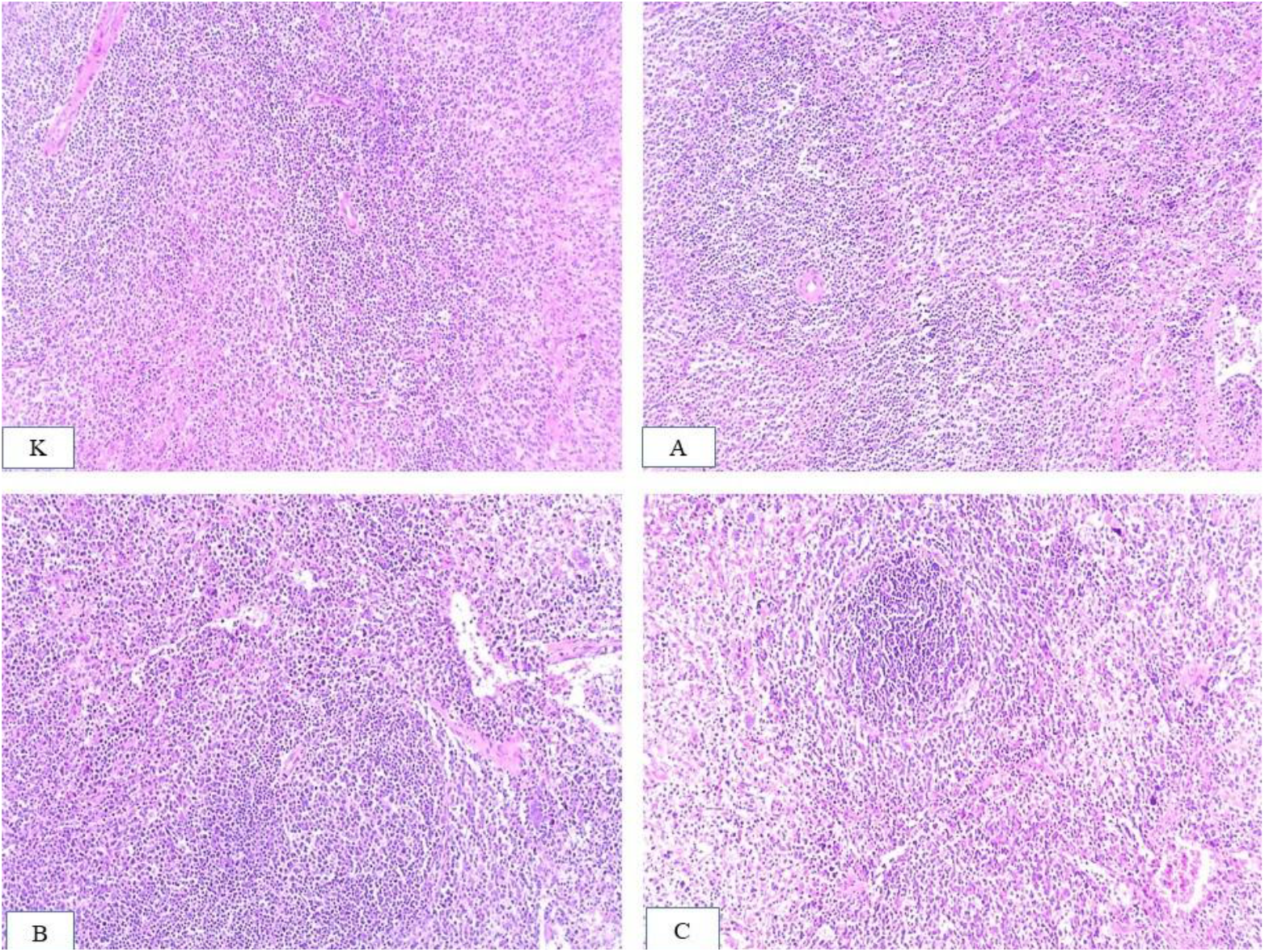
Normal spleen histology was observed in the K group, and mild inflammation/hemorrhage was observed in all treatment groups. HE 200x.

In the post hoc test of histological scores, as shown in Table 4, the liver damage scores in all treatment groups A, B, and C were significantly different compared to the control group, while the liver damage scores among treatment groups A, B, and C were not significantly different. The histological condition of the liver is linear with the increase in SGOT and SGPT levels on the 14th day of treatment (just before the termination of the rats). The increase in SGOT-SGPT levels and the highest histological liver damage scores were sequentially found in groups C, B, and A.

**Table 4.** Post hoc test of histologic score.

| Groups | *p*-value |  |
| --- | --- | --- |
|  | Liver Histology | Renal Histology |
| K-A | 0.138 | 0.523 |
| K-B | 0.019 | 0.241 |
| K-C | 0.005* | 0.005* |
| A-B | 0.269 | 0.575 |
| A-C | 0.093 | 0.019* |
| B-C | 0.523 | 0.056 |

\*statistically significant results

Meanwhile, in kidney histology, only the damage score in treatment group C was significantly different from the control group. The score of treatment group C also significantly differed from the score of treatment group A. The histological condition of the kidney is linear, with the increase in urea levels on the 14th day of treatment (just before the termination of the rats), but not for creatinine levels. The increase in urea levels and the highest kidney histological damage scores were found in treatment group C, followed by treatment groups B and A.

According to our research, intravenously administering Sprague Dawley rats a single-dose infusion solution containing NO, magnesium, and hydrogen nanobubbles does not result in mortality (LD_50_ = 0). The increase in urea and SGOT-SGPT levels is still within normal limits. Along with damage seen in the liver and kidney histology, this study additionally revealed an increase in SGOT-SGPT and urea levels, which were linearly correlated with a decrease in HNB dosage and an increase in NO dosage in the test preparation.

Our findings demonstrate that a single intravenous dose of the NO, Mg, and HNB mixture does not significantly harm test animals. The results of this study support earlier investigations into the effectiveness and safety of HNB in rat models of hypercholesterolemia (Hernowo et al., 2024). In addition, HNB demonstrated the potential to suppress adipogenesis, inflammatory response, and oxidative stress at the cellular and tissue levels (Xiao & Miwa, 2021). Meanwhile, Mg^2+^ with the same dosage in each treatment group did not significantly impact the results of this study.

The role of NO in this study still has to be investigated further. The function of NO bioactivities is multifaceted and complex. Inhaled NO is the most prevalent type of NO utilized in medical therapy, and it offers significant benefits, especially for heart and lung diseases (Signori et al., 2022). There are still a few studies on NO in injectable formulations. Previous animal studies have shown that infusing a saturated NO solution during hypoxic pulmonary hypertension can inhibit pulmonary vasoconstriction, most likely resulting from improved coronary circulation and cardiac inotropy (Pirrone et al., 2007). Meanwhile, another study found that eNOS-derived NO protects against liver and kidney illness, but iNOS-derived NO is detrimental (Hollenberg, 2006; Iwakiri & Kim, 2015).

In conclusion, the solution containing NO, Mg^2+^, and HNB has no serious adverse effects, which renders it safe for intravenous administration. To evaluate the possible long-term harmful effects, further research is highly recommended.

## Acknowledgment

This research was funded by Indonesia Molecule Institute in collaboration with Faculty of Medicine Universitas Jenderal Soedirman.

